# *In vivo*, regulated reconstitution of spindle checkpoint arrest and silencing through chemical-induced dimerisation

**DOI:** 10.1101/406983

**Authors:** Priya Amin, Sadhbh Soper Ní Chafraidh, Ioanna Leontiou, Kevin G. Hardwick

## Abstract

Chemical-induced dimerisation (CID) uses small molecules to control specific protein-protein interactions. Here, we employ CID dependent on the plant hormone abscisic acid (ABA) to reconstitute spindle checkpoint signalling in fission yeast. The spindle checkpoint signal usually originates at unattached or inappropriately attached kinetochores. These are complex, multi-protein structures with several important functions. To bypass kinetochore complexity, we take a reductionist approach to study checkpoint signalling. We generate a synthetic checkpoint arrest ectopically by inducing hetero-dimerisation of the checkpoint proteins Mph1^Mps1^ and Spc7^KNL1^. These proteins are engineered such that they can’t localise to kinetochores, and only form a complex in the presence of ABA. Using this novel assay we are able to checkpoint arrest a synchronous population of cells within 30 minutes of ABA addition. This assay allows for detailed genetic dissection of checkpoint activation and importantly it also provides a valuable tool for studying checkpoint silencing. To analyse silencing of the checkpoint and the ensuing mitotic exit, we simply wash-out the ABA from arrested cells. We show here that silencing is critically-dependent on PP1^Dis2^ recruitment to Mph1^Mps1^-Spc7^KNL1^ signalling platforms.

## Introduction

Spindle checkpoint signalling was initially reconstituted in *Xenopus* egg extracts (Kulukian et al., 2009; Minshull et al., 1994) and most recently using recombinant complexes of human checkpoint proteins (Faesen et al., 2017). Major advantages of such *in vitro* assays is that complex systems can be simplified through biochemical fractionation and manipulated through immunodepletion. They also enable the regulated addition of specific components, where the timing, concentration and activity of these can all be varied.

In parallel, yeast genetics drove identification of most of the molecular components of this pathway, the Mad and Bub proteins (Hoyt et al., 1991; Li and Murray, 1991), and their Cdc20 effector (Hwang et al., 1998; Kim et al., 1998). This combination of yeast genetics and *in vitro* reconstitution has proven invaluable when dissecting the molecular mechanism of action of spindle checkpoint signals and inhibition of the downstream effector Cdc20-APC/C (London and Biggins, 2014; Musacchio, 2015).

Here we have employed a hybrid approach, using yeast genetics and partial reconstitution of the pathway *in vivo*. We use synthetic biology to re-wire and simplify the upstream part of the checkpoint signalling pathway and chemical induced dimerisation (CID) to add an extra level of regulation that can be easily controlled experimentally in intact cells. Employing this strategy, we:

1) simplify the system, through regulated, ectopic activation of the spindle checkpoint, enabling kinetochore-independent studies.

2) use yeast genetics to enable rapid iterative analyses.

3) employ synthetic biology and CID to generate specific complexes in an experimentally-controlled fashion.

4) use abscisic acid (ABA) addition and wash-out to provide tight temporal control of the initiation and termination of checkpoint signalling.

More specifically, we generate a synthetic checkpoint arrest ectopically by inducing hetero-dimerisation of the checkpoint proteins Mph1^Mps1^ and Spc7^KNL1^ in fission yeast. This leads to checkpoint arrest in a synchronous population of cells within 30 minutes of addition of the plant phytohormone abscisic acid (ABA). As expected this checkpoint response requires the downstream Mad and Bub factors. To analyse silencing of the checkpoint, we simply wash-out the ABA from arrested cells and analyse mitotic exit. We find that the kinetics of release is critically-dependent on PP1^Dis2^ recruitment to the Mph1^Mps1^-Spc7^KNL1^ signalling platform.

## Results

We previously published a synthetic checkpoint arrest assay (SynCheck) where we activated the spindle checkpoint in fission yeast using heterodimers of TetR-Spc7 and TetR-Mph1 kinase (Yuan et al., 2017). However, in those experiments, dimerisation was constitutive, being driven by formation of Tet repressor dimers (TetR) and thus checkpoint signalling was challenging to regulate both in terms of initiation and termination. We controlled checkpoint arrest at the transcriptional level, using an *nmt* promoter to drive expression of the TetR-Mph1 fusion protein. Unfortunately the fission yeast *nmt1* promoter requires induction in media lacking thiamine for several hours. As a consequence, the peak of arrest was observed ∼14 hours after induction and wasn’t as synchronous as one would wish. Here, to improve both timing and control, we have modified our approach by employing chemical-induced dimerisation (CID) to give us tight temporal control over the initiation and termination of checkpoint signalling.

### Generation of SynCheckABA

Following the strategy of Crabtree and colleagues (Liang et al., 2011) we fused the PYL domain (residues 33 to 209) of the ABA receptor after the N-terminal 666 amino acids of fission yeast Spc7^KNL1^. By deleting the C-terminal half of Spc7^KNL1^ this protein is unable to be targeted to kinetochores, as it lacks the Mis12-interacting region (Petrovic et al., 2016; Petrovic et al., 2014). This fusion protein was expressed from the constitutive adh21-promoter (Tanaka et al., 2009). The ABI domain (residues 126-423) of ABI1 was fused to the C-terminus of the Mph1 spindle checkpoint kinase. We also deleted the first 301 amino-acids of Mph1 to prevent it going to kinetochores (Heinrich et al., 2012). This Mph1-ABI fusion protein was expressed from the adh41-promoter (Tanaka et al., 2009). In the presence of abscisic acid the PYL and ABI domains are sufficient to form a tight complex (Miyazono et al., 2009), thus forming Mph1^ABI^- Spc^7PYL^ complexes (Fig. 1A). We combined these constructs in a strain that also had the *cdc25-22* mutation, enabling synchronisation in G2, Bub1-GFP and mCherry-Atb2 to label microtubules.

**Figure 1.**
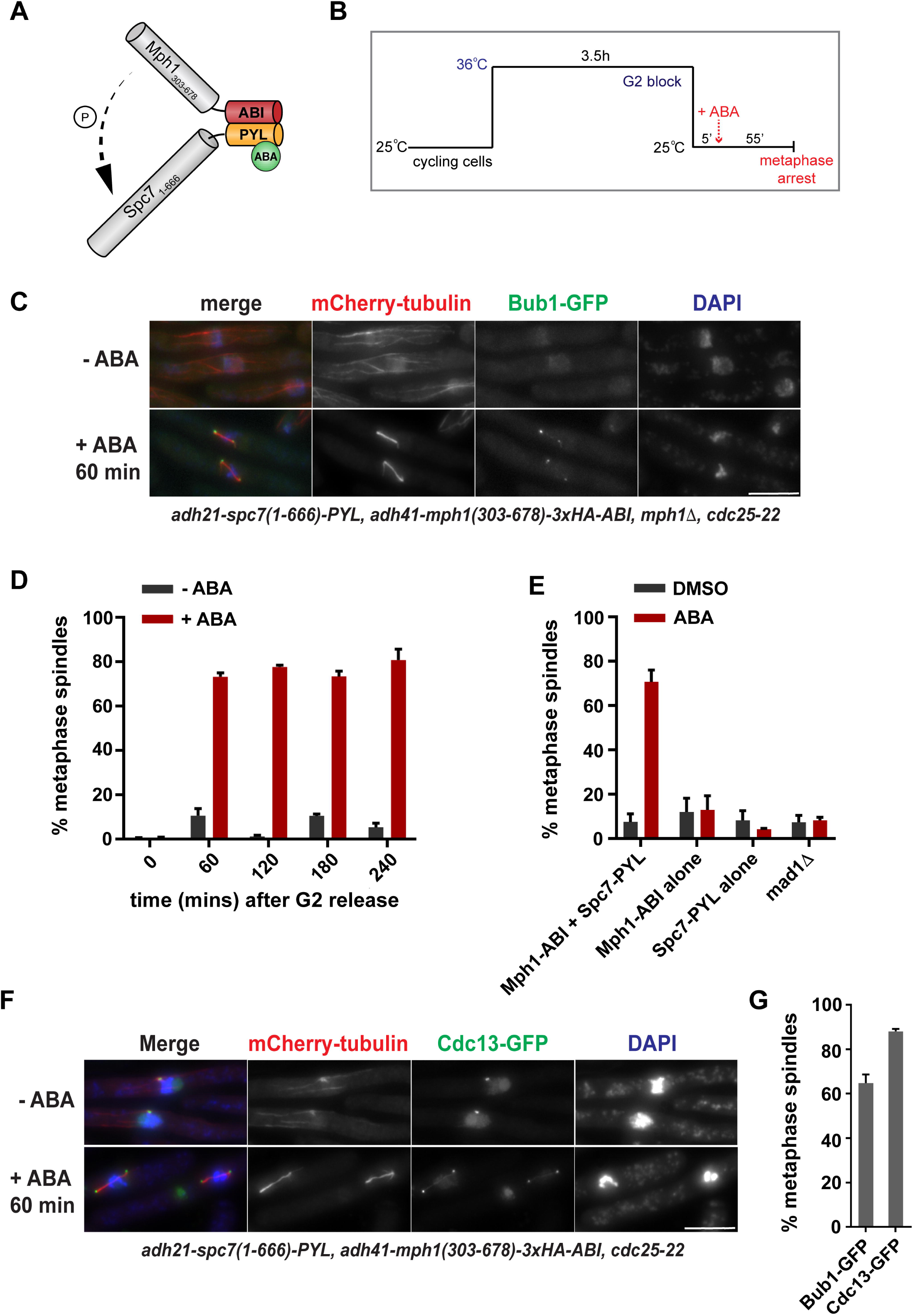
Rapid induction of spindle checkpoint arrest using abscisic acid for chemical-induced dimerisation of Mph1^Mps1^-Spc7^KNL1^. (A) Schematic representation of the Mph1^Mps1^-Spc7^KNL1^ heterodimer induced by ABA addition. (B) Work flow of the pre-synchronisation in G2 (cdc25-22), followed by release into mitosis at 25° C and then induction of checkpoint arrest through the addition of ABA. (C) Fixed cell images taken of the arrested ABA-induced strain 60 minutes after ABA addition. Microtubules are seen in red mCherry-Atb2), the checkpoint protein in green (Bub1-GFP) and chromatin is stained with DAPI. Scale bar is 10 μm. (D) Quantitation of cultures (+/- ABA addition) through a 4 hour time course after release from G2. Samples were fixed every 60 minutes and scored as metaphase arrested if they had short metaphase spindles and a single mass of condensed chromatin. >100 cells were analysed per strain at each time point. This experiment was repeated 3 times. Data plotted as mean +/- SD. (E) Quantitation of the strains indicated at the 60 minute time point after release from the G2 block (ABA added 5 mins after release). *mad1Δ* is the Mph1-ABI Spc7-PYL strain with *mad1* deleted. Cells were scored as metaphase arrested if they had short metaphase spindles and a single mass of condensed chromatin. 100 cells were analysed per strain at each time point. This experiment was repeated 3 times for each strain. Data plotted as mean +/- SD. (F) Fixed cell images taken of the SynCheckABA strain with Cdc13-GFP at spindle poles bodies 60 minutes after ABA addition. Microtubules are seen in red (mCherry-Atb2 is labeled fission yeast tubulin), cyclin B in green (Cdc13-GFP) and chromatin is stained with DAPI. Scale bar is 10 μm. (G) Quantitation comparing an ABA-induced metaphase arrest at 60 minutes between an Mph1-ABI Spc7-PYL strain containing Bub1-GFP and another Cdc13-GFP. This experiment was repeated 2 times. Data plotted as mean +/- SD.

### Inducing Spc7-Mph1 heterodimers to trigger a metaphase arrest

Cells were synchronised in G2 using a temperature-sensitive *cdc25-22* mutant which blocks cells in G2 after 3.5h at 36°C. When cells were shifted to 25°C, they ‘released’ from the block enabling progression through the cell cycle. After 5 minutes, ABA was added to activate the spindle checkpoint through the formation of Spc7-PYL and Mph1-ABI heterodimers (Fig. 1B). We observed that over 70% of cells had short metaphase spindles 60 minutes after ABA addition to the synchronous population of cells (Fig. 1C&D). The metaphase arrest could be sustained for at least 4 hours (Fig. 1D). We tested a range of ABA concentrations (0-500μM) and found that 250μM was optimal for reproducible, robust arrests (Fig. S1A). The ABA can be added later (eg. 20 mins after *cdc25* release) and cells arrest with similar efficiency to that observed after anti-microtubule drug treatment (carbendazim, CBZ, see Fig. S1B). Without pre-synchronisation in G2, the mitotic index increases over time and reaches a peak four hours after ABA addition (Fig. S1C). In our previous SynCheck studies cells arrested for several hours but then died [10]. We wanted to determine whether the ABA arrest also had a significant affect on cell viability or whether our ability to release this arrest (through ABA wash-out) meant that viability was maintained. After ABA treatment we found a gradual drop in cell viability (see Fig. 2E), which was similar to that observed upon anti-microtubule drug treatment (data not shown). In the arrested cells, we observed Bub1 enrichment at the spindle poles (Fig. 1C). This is consistent with our previous SynCheck assay where movement of all spindle checkpoint proteins to spindle poles was reported to be Mad1-Cut7 kinesin driven (Yuan et al., 2017). As expected, deleting the first N-terminal coiled coil (136 amino acids) of Mad1, required for its interaction with the Cut7 (Akera et al., 2015), prevented Bub1 accumulation at spindle poles. This de-localisation of checkpoint proteins from spindle poles did not affect the efficiency of the arrest (Fig. S1D), as was found in SynCheck (Yuan et al., 2017).

**Figure 2.**
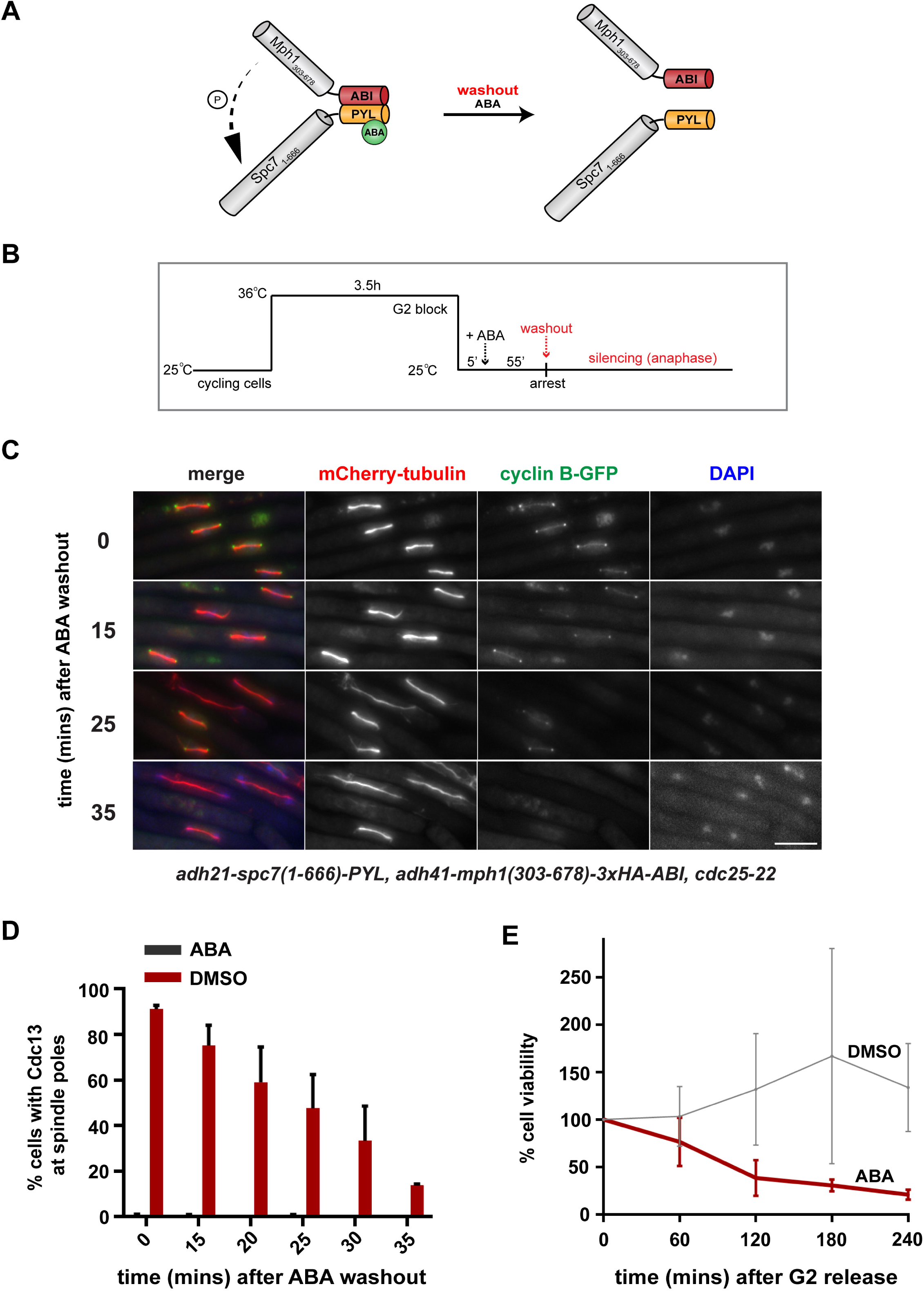
Silencing of spindle checkpoint signalling after abscisic acid wash-out. (A) Schematic representation of the dissociation of Mph1^Mps1^-Spc7^KNL1^ heterodimers after ABA wash-out. (B) Silencing work flow: pre-synchronisation in G2 (*cdc25-22*), induction of checkpoint arrest through the addition of ABA, and subsequent wash-out of ABA 60 minutes later. (C) Fixed cell images taken of the arrested SynCheckABA strain at 0, 15, 25 and 35 minutes after ABA wash-out. Microtubules are seen in red (mCherry-Atb2), cyclin Bin green (Cdc13-GFP) and chromatin is stained with DAPI. Scale bar is 10 μm. See Fig. S2B for an alternatively coloured version of similar images. (D) Quantitation of Cdc13-GFP at spindle pole bodies in the SynCheckABA cultures (+ ABA/DMSO). Samples were fixed and scored for the presence of Cdc13 at spindle pole bodies. The +DMSO control didn’t arrest in metaphase. >150 cells were analysed per strain at each time point. This experiment was repeated 3 times. Data are plotted as mean +/- SD. (E) The viability of SynCheckABA arrested strains was determined by plating cells 0, 60, 120, 180 and 240 minutes after release from a G2 block where DMSO or ABA was added 5 minutes after release from the G2 block. Cell viability over time was plotted as a percentage relative to that at time zero. Cells were plated in triplicate. The experiment was repeated 3 times and data are plotted as mean +/- SD.

ABA-induced metaphase arrest is dependent on hetero-dimerisation of Spc7-PYL and Mph1-ABI. Strains lacking either the Mph1-ABI component, or the Spc7-PYL component, failed to arrest in the presence of ABA (Fig. 1E). The ABA-induced arrest is checkpoint-dependent, as deleting the downstream checkpoint protein Mad1 abolished the arrest (Fig. 1E). In these constructs Spc7 and Mph1 lack their kinetochore-binding domains, making initiation of this arrest ectopic and independent of the complexities of the kinetochore. The Mph1-ABI, Spc7-PYL strain used above lacks endogenous *mph1*, which prevents all Mad/Bub checkpoint proteins from targeting to kinetochores (Heinrich et al., 2012). As an additional measure, to confirm kinetochore independence, we employed a strain containing the *spc7-12A* MELT mutant allele (Mora-Santos et al., 2016; Yamagishi et al., 2012). This mutant Spc7 kinetochore component cannot be phosphorylated by Mph1, preventing recruitment of Bub3-Bub1 and thereby Mad1-Mad2 complexes to kinetochores. The *spc7-12A* mutant arrested with very similar efficiency to *spc7+* cells under ABA control (Fig. S1F), arguing that the Spc7wt-PYL Mph1-ABI heterodimer does not need to be aided by endogenous kinetochore-based checkpoint signalling to generate a checkpoint arrest. Importantly, an *spc7-12A-PYL* fusion protein was unable to generate an arrest in combination with Mph1-ABI, demonstrating that the ectopic signaling scaffold does need to be phosphorylated on conserved Spc7 MELT motifs to recruit Bub3-Bub1 complexes for active signaling (Fig. S1G).

Critical outputs of checkpoint action are the stabilisation of cyclin B and securin. Using a modified strain we analysed cyclin B (Cdc13) levels in the ABA-induced arrest. Fig. 1F shows that Cdc13-GFP accumulated on short metaphase spindles and was enriched at mitotic spindle poles, as expected. As a technical aside, we have found that different tags can affect the efficiency of the ABA-induced arrest. For example, this Cdc13-GFP strain reproducibly arrests more efficiently than the strain containing Bub1-GFP (Fig. 1G). This is likely due to a partial loss of function when C-terminally tagging the Bub1 checkpoint protein. The Cdc13-GFP strain also contains the endogenous wild-type *Mph1* gene, but we find that this does not significantly impact the efficiency of arrest (see Fig. S1E).

Thus we have reconstituted a robust, kinetochore-independent checkpoint arrest that can be initiated very simply *in vivo* through ABA addition to culture media. This works efficiently in both minimal (PMG) and rich (YES) fission yeast growth media. Hereafter, we refer to this assay as SynCheckABA.

### A novel spindle checkpoint silencing assay

A significant advantage of SynCheckABA is the ability to reverse the effects of abscisic acid by simply washing cells with fresh media lacking ABA and thereby releasing them from metaphase arrest (Fig. 2A/B). We can use this to study spindle checkpoint silencing, which has proven to be technically challenging in the past. Fig. 2C/D demonstrates that when we wash-out the ABA we observe rapid cyclin degradation and spindle elongation (see also Fig. S2A).

### Regulation of spindle checkpoint silencing

Previous work has shown that PP1^Dis2^ is a key spindle checkpoint silencing factor in yeasts (Meadows et al., 2011; Pinsky et al., 2009; Vanoosthuyse and Hardwick, 2009). The N-terminus of Spc7^KNL1^ has two conserved motifs (SILK and RRVSF, also referred to as the A and B motifs) mediating stable PP1^Dis2^ association (Fig. 3A). Mutation of both binding sites leads to a lethal metaphase block in *S. cerevisiae* and in *S. pombe* (Meadows et al., 2011; Rosenberg et al., 2011). There are additional kinetochore binding sites for PP1^Dis2^ such as Klp5 and Klp6 (Meadows et al., 2011) and these are relevant to checkpoint silencing, although binding to Spc7^KNL1^ appears to be the major player. In human cells similar motifs are found at the N-terminus of KNL1 and PP1 binding is regulated by Aurora B activity as this kinase can directly phosphorylate the B motif disrupting PP1 association (Liu et al., 2010).

**Figure 3.**
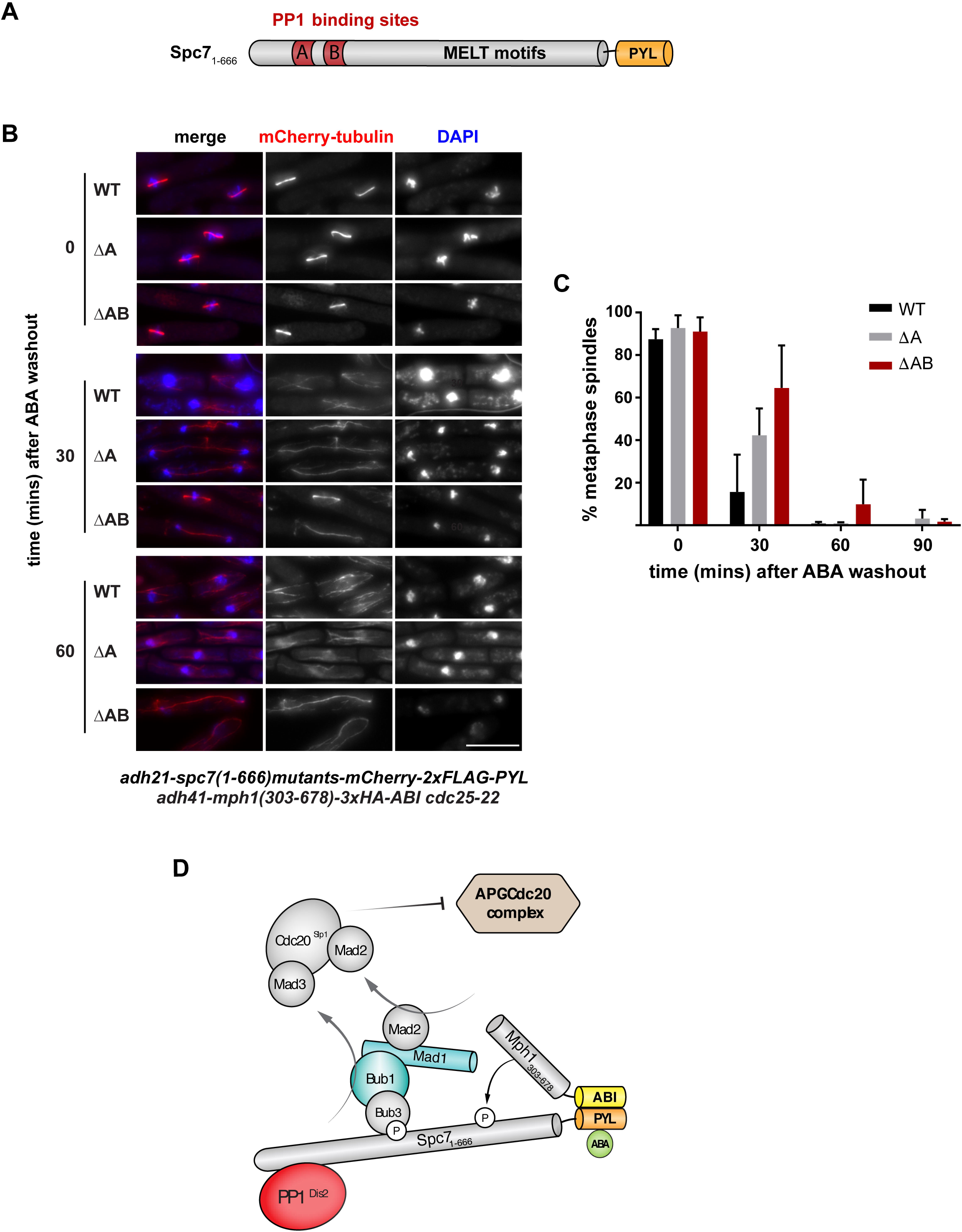
Checkpoint silencing in SynCheckABA is dramatically slowed when the Spc7^KNL1^ binding sites for PP1^Dis2^ are deleted. (A) Schematic of Spc7^KNL1^ indicating the N-terminal PP1 binding motifs (A motif: SILK and B motif: RRVSF). The MELT motifs form binding sites for Bub3-Bub1 complexes once they have been phosphorylated by Mph1 kinase. (B) Images of cells expressing wild-type Spc7_1-666_ or mutants with deletion of the A motif or both the A and B motifs. Time points were analysed at the time of ABA washout (time zero) and 30 and 60 minutes post-wash. Scale bar is 10 μm. See Fig. S3 for non red-green colour scheme. (C) Quantitation of this release from checkpoint arrest in the strains expressing wild-type Spc7_1-666_or mutants with deletion of the A motif or deletion of both the A and B motifs. This experiment was repeated three times. >100 cells were analysed per strain at each time point. Data are plotted as mean +/- SD. (D) Schematic model of SynCheckABA: activating (Mph1) and silencing (PP1) factors bind nearby on the Spc7 scaffold. The balance of their activities will determine how much MCC is generated and thus whether anaphase onset is inhibited.

Employing SynCheckABA, we tested mutations of the A and B motifs at the N-terminus of Spc7^KNL1^ and removal of the Klp5 and Klp6 kinesins. For the experiments below (Figs. 3 and 4), all strains contained endogenous, wild-type *Mph1* kinase and thus are able to recruit checkpoint proteins to their kinetochores. This includes the Mph1 and Bub1 kinases which are also thought to also have ‘error correction’ functions. Thus silencing likely needs to take place not only at the ectopic Mph1^Mps1^-Spc7^KNL1^ signalling scaffold, but also at kinetochores. Strains were pre-synchronised in G2 using *cdc25*, released and arrested at metaphase using ABA, and then washed to terminate checkpoint signalling. Progression through anaphase was then scored through the analysis of spindle elongation and/or cyclin B degradation (using Cdc13-GFP) over a 90 minute time-course. Mutation of the A-motif delayed spindle elongation by 30 minutes and the A/B double mutant was delayed even more profoundly (Fig. 3B/C). This argues that PP1 activity on or near the Spc7 protein (previously phosphorylated by Mph1 kinase) is a limiting factor in checkpoint silencing. This system will prove useful for dissecting the regulation of PP1 binding to Spc7 in more detail, and for analysis of putative regulators of PP1 activity.

**Figure 4.**
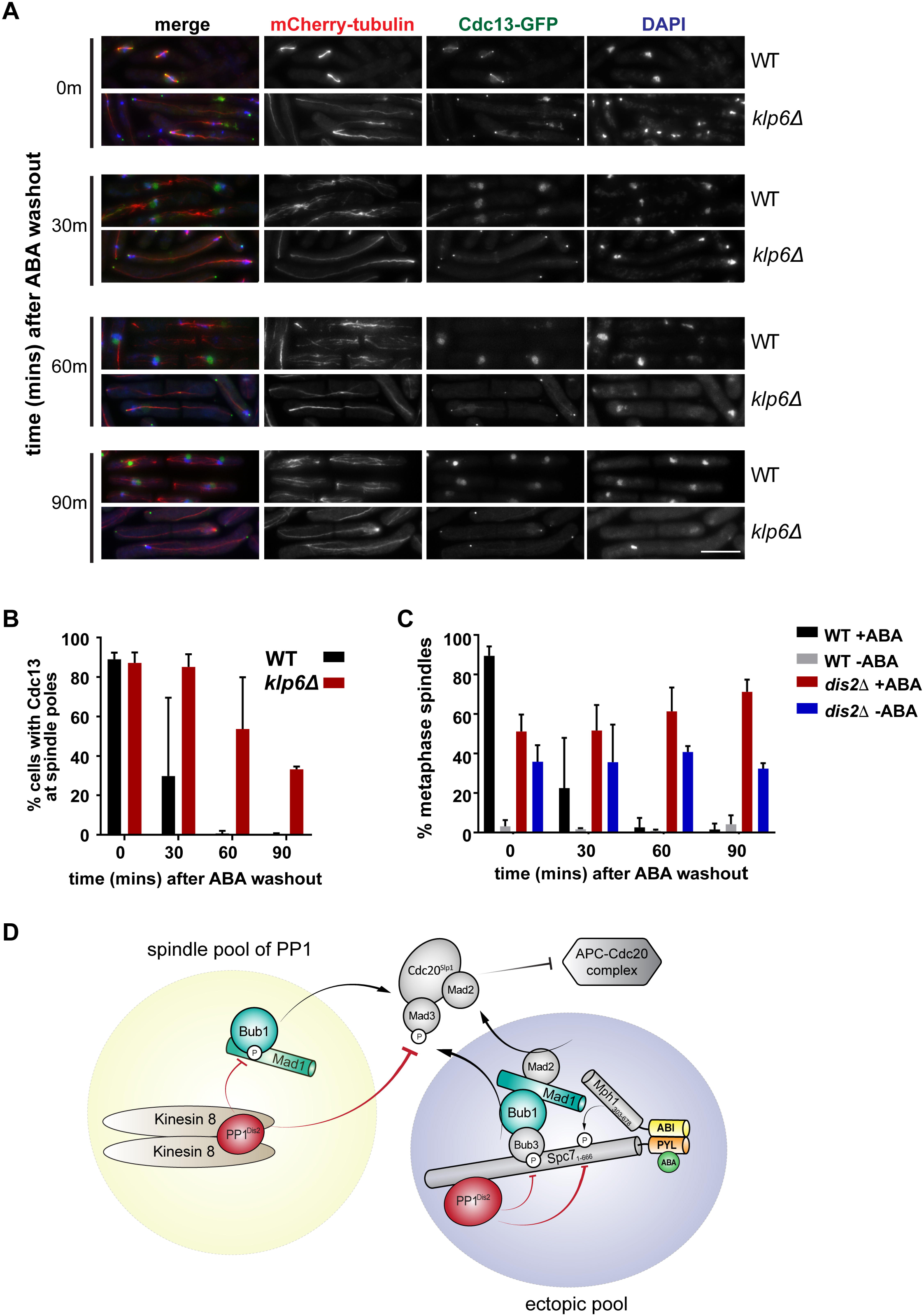
Checkpoint silencing in SynCheckABA is also slowed when other recruitment sites for PP1^Dis2^ are removed from spindles. (A) Deletion of the kinesin 8, Klp6, leads to reduced silencing efficiency. Images of cells with and without Klp6 deleted are shown after ABA washout (time zero) and 30, 60 and 90 minutes post-wash. Microtubules are seen in red (mCherry-Atb2), cyclin Bin green (Cdc13-GFP) and chromatin is stained with DAPI. Scale bar is 10μm. (B) Quantitation of this release from checkpoint arrest in strains with and without Klp6. Cells were scored as arrested if Cdc13-GFP was enriched at spindle poles. This experiment was repeated three times. >100 cells were analysed per strain at each time point. Data plotted as mean +/- SD. (C) *dis2*- mutants have profound silencing defects. Quantitation of the release from the checkpoint arrest is shown for wild-type and *dis2*- cells (+ABA/DMSO). Cells were scored as metaphase arrested if they had short metaphase spindles and a single mass of condensed chromatin. DMSO controls illustrate that *dis2*- cells are generally sick, but that ABA addition induces the SynCheckABA, resulting in elevated levels of metaphase arrested cells. This arrest persists for >60m after ABA washout as *dis2*- cells struggle to silence the checkpoint. This experiment was repeated three times. >200 cells were analysed per strain at each time point. Data plotted as mean +/- SD. See Fig. S4 for fixed cell images from this timecourse. (D) General model of relevant PP1-dependent silencing pathways. This schematic describes two pools of PP1^Dis2^ one that is recruited to the ectopic Spc7-Mph1 signalling scaffold, via the A and B motifs on Spc7, and a second pool that is more generally recruited to the spindle through interaction with kinesin 8 (Klp6). These two pools act together to inhibit MCC-APC/C assembly and thereby enable checkpoint silencing and mitotic exit.

Mutation of fission yeast kinesin 8 (either Klp5 or Klp6) leads to a stabilisation of microtubules, aberrant chromosome movements and long metaphase spindles (Gergely et al., 2016; Klemm et al., 2018; Meadows et al., 2011; West et al., 2002). In these mutants, checkpoint silencing defects can’t simply be analysed through spindle elongation. Instead we imaged Cdc13-GFP and used the decrease in the number of cells with cyclin B enriched at their spindle poles as a measure of checkpoint silencing. Fig. 4A/B demonstrate that mutation of Klp6 significantly reduces the efficiency of silencing and cyclin B degradation. Finally, we analysed the silencing defect upon deletion of the PP1^Dis2^ phosphatase (*dis2Δ*). In the *dis2Δ* strain, which is rather sick, checkpoint silencing was extremely defective with no significant drop in Cdc13-GFP levels over the 90 minute time course (Fig. 4C). It should be noted that these *dis2Δ* strains display significant mitotic delays even in the absence of ABA addition, presumably because the lack of this mitotic phosphatase leads to pleiotropic mitotic defects (see Fig. S4 for images of these cells).

Thus, SynCheckABA neatly recapitulates the balance of opposing kinase and phosphatase activities between Mph1^Mps1^ dependent checkpoint activation and PP1-driven checkpoint silencing, on Spc7^KNL1^ and Kinesin 8 dependent pathways (see general model in Fig. 4D).

## Discussion

Here we have employed chemical-induced dimerisation to generate a rapid, controlled spindle checkpoint arrest: addition of abscisic acid (ABA) to SynCheckABA strains induces the heterodimerisation of Mph1^Mps1^-ABI and Spc7^KNL1^-PYL fusion proteins and this is sufficient to generate an activated signalling scaffold and metaphase arrest within minutes. Like our original SynCheck assay, which was driven by constitutive TetR homodimers (Yuan et al., 2017), this arrest acts independent of spindle checkpoint signalling at endogenous kinetochores, but is dependent on downstream checkpoint components such as Mad1.

A significant advantage of SynCheckABA is that we can wash out the ABA and study the kinetics and mechanism of spindle checkpoint silencing. This was not possible with the original SynCheck strain as we were unable to control TetR dimerisation and thus unable to dissociate the Mph1^Mps1^-TetR-Spc7^KNL1^-TetR signalling scaffold.

Using this new assay we have confirmed that protein phosphatase 1 (PP1^Dis2^) is critical for silencing the Mph1^Mps1^- Spc7^KNL1^ scaffold (Figs. 3-4). PP1^Dis2^ binds to the N-terminus of Spc7^KNL1^, not far from the conserved MELT motifs that, once phosphorylated by Mph1^Mps1^, will bind Bub3-Bub1 complexes to initiate MCC generation (Shepperd et al., 2012). Thus Spc7^KNL1^ acts as the platform for both checkpoint activation and silencing and appears to be a major site of action of both checkpoint activation kinases and silencing phosphatases (Meadows et al., 2011). It is important to note that not all aspects of silencing are recapitulated in our ectopic assay, as some of these will relate to specific kinetochore processes that will not be captured here.

Kinesin 8 is also confirmed as a PP1^Dis2^ recruitment site relevant for checkpoint silencing in SynCheckABA. The phenotypes of the *klp6Δ* mutant suggests that targeting of PP1 to spindle microtubules and kinetochores is also relevant to mitotic exit from an ABA-induced arrest, even though the arrest is initiated away from the kinetochore (see Fig. 4D).

### Advantages of SynCheckABA, over other forms of reconstitution

We believe that all forms of spindle checkpoint reconstitution are useful for mechanistic dissection of this dynamic signalling pathway, whether this be *in vitro* within cytoplasmic extracts (Minshull et al., 1994), *in vitro* with purified recombinant proteins (Faesen et al., 2017) or *in vivo* with synthetically re-wired and simplified signalling pathways (SynCheckABA). The advantages of the latter system are:

1) the signalling pathway downstream of Spc7^KNL1^ and the downstream effectors are present at normal physiological levels and there are simple, quantitative physiological read-outs (cyclin B degradation, sister chromatid separation and/or anaphase spindle elongation).

2) checkpoint arrest is induced in the absence of additional stresses: simple addition of abscisic acid (low toxicity) to the growth media is sufficient for checkpoint activation. There is no need for a cold-shock (to depolymerise tubulin, *nda3* arrest), heat-shock (to perturb temperature-sensitive kinetochore mutants), or overexpression of checkpoint activators.

3) The PYL and ABI domains have limited cross-reaction in yeast as they are derived from plant proteins. Although we haven’t compared them directly, we believe that abscisic acid has certain advantages over the use of rapamycin, a very popular CID. To use rapamycin in fission yeast one needs to engineer strains to remove endogenous rapamycin-binding proteins, such as by deleting the *fkh1*+ gene which encodes a native FKBP12 domain (Ding et al., 2014).

Importantly, because ABA doesn’t bind tightly to the PYL domain, we can wash ABA out easily to initiate checkpoint silencing. In comparison, rapamycin is very difficult to wash-out making efficient release experiments unrealistic.

4) compared to *in vitro* studies with large, recombinant complexes, these fission yeast experiments are simple, cheap and fast. The system also enables rapid iterative studies, due to ease of further genetic manipulation in yeast.

5) importantly, we can easily test candidate regulators (eg. silencing factors) without needing to know what complexes they are part of, purifying them and worrying about their relevant concentration and post-translational modifications.

6) compared to our transcriptionally controlled SynCheck (which employs *nmt-tetR-Mph1*) the ABI-PYL system is less leaky, enabling sick strains such as the *dis2* mutant analysed in Fig. 4 to be constructed. Previously, we were unable to isolate *nmt-tetR-Mph1, dis2*- strains, due to leaky expression from the weak nmt81 promoter.

Our ongoing studies with SynCheckABA will enable a detailed mechanistic dissection of PP1-mediated spindle checkpoint silencing in fission yeast. We believe that ABA holds a lot of promise as an alternative CID to rapamycin and indeed that it has significant advantages.

## Acknowledgements

We would like to thank Patrick Heun and Eftychia Kyriacou for providing constructs containing the abscisic acid binding hetero-dimerisation domains of PYL and ABI; Jonathan Millar for the PP1 binding *spc7* mutants and the *klp5,6* mutants; and Ken Sawin for the mCherry-Atb2 strain. This work was supported by a Seed Award from the Wellcome Trust to K.G.H. (108105) and the Wellcome Centre for Cell Biology core grant (092076). P.A. was supported by the Medical Research Council (MR/K501293/1), S.S.N.C. by the Wellcome Trust (105258) and I.L. by the Darwin Trust of Edinburgh.

## Materials and Methods

### P_adh41_-mph1_303-678_-3xHA-ABI

Mph1_303-678_ was amplified from a pDONR 201 plasmid containing Mph1_303-678_ (Yuan et al., 2017). 3xHA was amplified from a plasmid from the Allshire lab containing codon optimised PYL-3xHA. ABI was amplified from a pMT_CID_ABI_VS_H vector from the Heun lab. These PCR fragments were Dpn1 treated and assembled into a Sma1-digested and antarctic phosphatase treated, gel purified pRad41 yeast expression vector by Gibson assembly.

### P_adh21_-spc7_1-666_-PYL

A yeast expression vector (pIY03 (Yuan et al., 2017)) was digested with Nhe1 and Xho1 and gel purified. The insert (mCherry-3xFlag-Spc7_1-666_) was used as a template to amplify Spc7_1-666_. PYL was amplified from a bVNI-221 vector from the Heun lab. The fragments were then assembled into the digested pIY03 vector backbone using Gibson assembly.

### P_adh21_-spc7_1-666_-mCherry-2xFLAG-PYL (PP1-binding site mutants)

Plasmids containing full length Spc7 PP1-binding mutants (ΔA: deletion of residues 136-150; ΔAB: deleting residues 136-150 and residues 331-345) (Millar lab) were used as templates to amplify mutant versions of Spc7_1-666_. NheI-NLS and PacI sites were introduced during amplification, allowing Spc7 constructs to be digested and ligated into digested pIY03-derived vector backbone which also contained a C-terminal mCherry-2Xflag-PYL tag.

### Fission yeast strains

These are listed in Supplementary Table 1.

### *cdc25-22* synchronisation

Cells were grown at 25°C for 1-2 days on YES (rich yeast media, with additional leucine, arginine, lysine, histidine and uracil) plates. They were then pre-cultured in 10 ml of liquid YES containing amino acid supplements at 25° over the day and inoculated into a larger culture of YES overnight. The following day, log phase cultures were shifted to 36°C for 3.5 hours to block in G2. After this, cultures were cooled quickly in iced water to rapidly shift them back to 25°C and release them from the G2 block.

### Synthetic arrest assay

Following a *cdc25-22* block, 250mM abscisic acid stock (Sigma Aldrich A1049) was added to cultures 5 minutes after release (20 minutes if comparing to a carbendazim arrest) to achieve a final concentration of 250uM (unless otherwise stated).

### Synthetic arrest assay wash-out

Following an ABA-induced synthetic arrest, the cells were washed 3 times (with 50 ml YES).

### Fixing cells and microscopy

1-1.5 ml of culture was centrifuged for 1 minute at 6000 RPM. The cell pellet was fixed in 200-500 ul 100% ice-cold methanol. To image cells, 8 ul of the cell-methanol suspension was added to a glass slide, when the methanol evaporated, 1-2 ul DAPI (0.4 ug/ml) was added to the sample and a glass cover slip was placed on top.

Cells were imaged immediately using a 100x oil immersion lens and a Zeiss Axiovert 200M microscope (Carl Zeiss Ltd.), equipped with a CoolSnap CCD camera (Photometrics) and Slidebook 5.0 software (3i, Intelligent Imaging Innovations, Inc.). Typical acquisition settings: 300 ms exposure (FITC & TRITC), 100 ms exposure DAPI, 2x binning, Z-series over 3 μm range in 0.5 μm steps (7 planes).

### Carbendazim arrest

Following a cdc25-22 block, 3.75mg/ml of carbendazim was added to cultures 20 minutes after release to achieve a final concentration of 100ug/ml.

### Cell viability assay

Following a synthetic arrest assay, cells from 1ml of culture were harvested by centrifugation at 6000 RPM for 1 minute and re-suspended in 1ml of distilled water. 10-fold serial dilutions were made in distilled water. 0.1 ml of cells diluted by factors of 100 and 1000 were plated in triplicate. Colony forming units per ml of culture was calculated and cell viability over time was plotted as a percentage relative to that at time zero.

